# 1-Methylnicotinamide is an immune regulatory metabolite in human ovarian cancer

**DOI:** 10.1101/2020.05.05.077990

**Authors:** Marisa K. Kilgour, Sarah MacPherson, Lauren Zacharias, Sarah Keyes, Brenna Pauly, Bertrand Allard, Julian Smazynski, Peter H. Watson, John Stagg, Brad H. Nelson, Ralph J. DeBerardinis, Phineas T. Hamilton, Julian J. Lum

## Abstract

Immune regulatory metabolites are key features of the tumor microenvironment (TME), yet with a few notable exceptions, their identities remain largely unknown. We uncovered the immune regulatory metabolic states and metabolomes of sorted tumor and stromal, CD4+, and CD8+ cells from the tumor and ascites of patients with high-grade serous ovarian cancer (HGSC) using high-dimensional flow cytometry and metabolomics supplemented with single cell RNA sequencing. Flow cytometry revealed that tumor cells show a consistently greater uptake of glucose than T cells, but similar mitochondrial activity. Cells within the ascites and tumor had pervasive metabolite differences, with a striking enrichment in 1-methylnicotinamide (MNA) in T cells infiltrating the tumor compared to ascites. Despite the elevated levels of MNA in T cells, the expression of nicotinamide N-methyltransferase, the gene encoding the enzyme that catalyses the transfer of a methyl group from S-adenosylmethionine to nicotinamide, was restricted to fibroblasts and tumor cells. Treatment of T cells with MNA resulted in an increase in T cell-mediated secretion of the tumor promoting cytokine tumor necrosis factor alpha. Thus, the TME-derived metabolite MNA contributes to an alternative and non-cell autonomous mechanism of immune modulation of T cells in HGSC. Collectively, uncovering the tumor-T cell metabolome may reveal metabolic vulnerabilities that can be exploited using T cell-based immunotherapies to treat human cancer.

---

Tumor-derived metabolites can have profound suppressive effects on anti-tumor immunity, with increasing evidence that they can also function as key drivers of disease progression^1,2^. Beyond the Warburg effect, recent work has begun to characterize the metabolic states of tumor cells and their relationship to the immunological state of the TME. Studies in murine models have helped uncover the role of metabolites such as (R)-2-hydroxyglutarate^3^, BH4^4^ and methylglyoxal5 as well as pathways including glutamine metabolism^6^, oxidative metabolism^7^, and glucose metabolism^8^ that impact T cell function and antitumor immunity. Furthermore, studies in humans have elucidated key metabolic pathways in tumors, for example demonstrating that tumors can use lactate as fuel^9^. Despite this, the diversity and impact of specific metabolites on tumor-infiltrating lymphocytes (TILs) are largely unknown. To characterize this diversity and better understand how metabolites in the TME influence T cell function, a combined flow cytometry and mass-spectrometry approach was used to profile tumor and TIL from patients with HGSC. Using this approach two spatially distinct microenvironments were interrogated, the ascites^10^ and tumor, within the same patients to reveal potential reciprocal metabolic interactions between tumor cells and TIL.

The phenotypic and metabolic states of cells in the matched ascites and tumor environments from six patients with HGSC (Extended Data Table 1) were evaluated using high-dimensional flow cytometry to synchronously quantify glucose uptake (2-(N-(7-Nitrobenz-2-oxa-1,3-diazol-4-yl)Amino)-2-Deoxyglucose, 2-NBDG) and mitochondrial activity (MitoTracker Deep Red)^5,11,12^ alongside canonical markers to distinguish immune and tumor cell populations (Extended Data Table 2, Extended Data Fig. 1a). This revealed high levels of glucose uptake in tumor cells relative to T cells in both the ascites and tumor, but more modest differences in mitochondrial activity. Tumor cells (CD45-EpCAM+) had on average 3-4 times the glucose uptake of T cells, whereas CD4+ T cells had on average 1.2 times the glucose uptake of CD8+ T cells, suggesting that TILs have different metabolic requirements even within the same TME (Fig. 1a). In contrast, the mitochondrial activity in tumor cells was similar to CD4+ T cells, and both had greater mitochondrial activity than CD8+ T cells (Fig. 1b). Collectively, these results reveal a metabolic hierarchy, with tumor cells more active than CD4+ T cells, and CD4+ T cells more metabolically active than CD8+ T cells. Despite these effects across cell types, there were no consistent differences in the metabolic states of CD4+ and CD8+ T cells, or their relative proportions, in the ascites compared to the tumor (Fig. 1c). Conversely, within the CD45-cell fraction, there was an increase in the proportion of EpCAM+ cells in the tumor compared to the ascites (Extended Data Fig. 1b). We also observed clear metabolic differences among EPCAM+ and EPCAM-cell fractions. EPCAM+ (tumor) cells had substantially greater glucose uptake and mitochondrial activity than EPCAM-cells, consistent with much higher metabolic activity in tumor cells than fibroblasts in the TME (Extended Data Fig. 1c, d).

**Figure 1.**
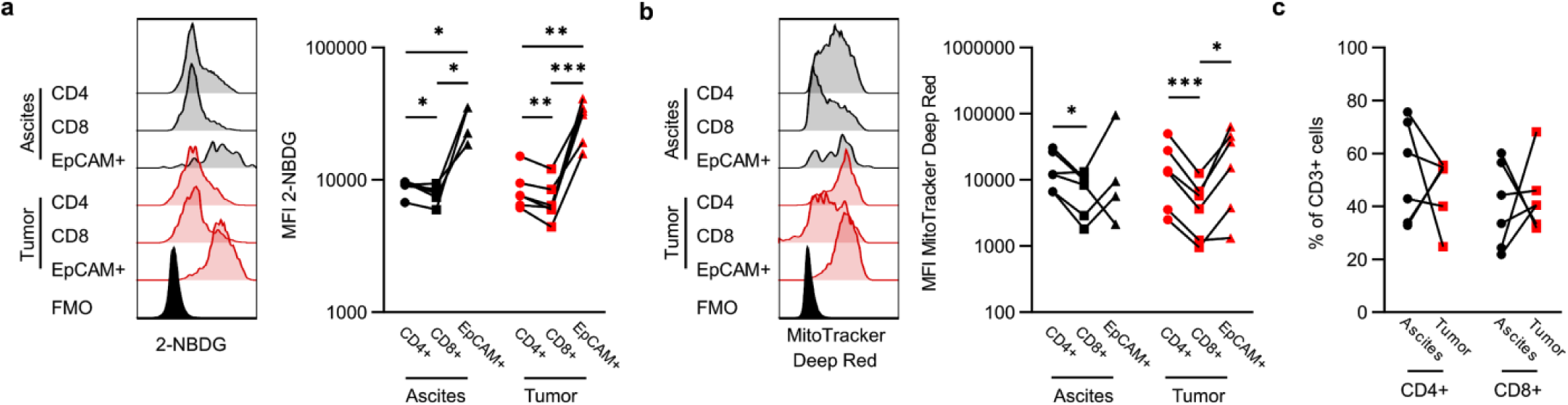
Tumor cells have greater glucose uptake but similar mitochondrial activity to T cells. **a, b**, Representative plot (left) and tabulated data (right) for median fluorescent intensity (MFI) of glucose uptake (2-NBDG) (**a**) and mitochondrial activity (MitoTracker Deep Red) (**b**) of CD4+ T cells, CD8+ T cells, and EpCAM+CD45-tumor cells from ascites and tumor. **c**, Proportion of CD4+ and CD8+ cells (of CD3+ T cells) within ascites and tumor. P-values determined by paired t-test (*p<0.05, **p<0.01, ***p<0.001) (**a**-**c**). Lines indicate matched patients (n=6). Fluorescence Minus One (FMO); Median Fluorescence Intensity (MFI).

Further analysis revealed other clear differences when considering more highly-resolved phenotypic states of T cells^13^. Indeed, activated (Extended Data Fig. 1e-g) and effector memory (Extended Data Fig. 1h, i) T cells were much more frequent (as a proportion of T cells) in the tumor than ascites. Similarly, resolving phenotypes by the expression of activation markers (PD1, CD25, CD137) revealed that while these populations showed some differences in metabolism (Extended Data Fig. 2a-e), no consistently significant metabolic differences were observed between naïve, effector, or memory cells (defined by CCR7 and CD45RO, Extended Data Fig. 2f-i). These results were confirmed through automated assignment of cell phenotypes using machine learning method^14^, which further revealed an abundant myeloid cell population (CD3-/CD4+) predominately in patient ascites that displayed the highest glucose uptake and mitochondrial activity of any identified cell type (Extended Data Fig 3). These results underscore strong metabolic differences across different cell types found in the ascites and tumors of HGSC patients.

A major challenge in understanding the metabolomic profiles of TIL has been the need to isolate samples of T cells of sufficient purity, quality and quantity from tumors. Recent studies have shown that flow cytometry based sorting and bead enrichment methods can cause alterations in cellular metabolite profiles^15–17^. To overcome this, we optimized a bead enrichment approach to isolate and separate TIL from surgically resected human ovarian cancers prior to analysis by liquid chromatography tandem mass spectrometry (LC-MS/MS) (See Methods; Extended Data Fig. 4a). To assess the overall impact of this protocol on metabolite changes, we compared the metabolite profiles of activated T cells following bead isolation to cells that did not undergo bead isolation but remained on ice, and found high correlation among methods (*r* = 0.77), as well as high reproducibility among technical replicates for this panel of 86 metabolites (see Extended Data Fig. 4b). These methods thus enabled accurate metabolite profiling in cells undergoing enrichment, to provide a first high-resolution platform for the identification of specific metabolites in HGSC thereby allowing deeper insight into cell-specific metabolic programs.

We applied this enrichment method to profile 99 metabolites in CD4+, CD8+, and CD45-cell fractions from the primary ascites and tumor of six patients with HGSC (Extended Data Fig. 4c). Profiling revealed strong metabolic separation of cell types within and across patients (Fig. 2a, Extended Data Fig. 5a). In particular, patient 70 had distinct metabolic profiles compared to other patients (Fig. 2b, Extended Data Fig. 5b), indicating the potential for substantial metabolic heterogeneity among patients. Notably, patient 70 had a smaller total volume of ascites collected (80 mL) compared to the other patients (1.2-2 L) (Extended Data Table 1). Controlling for inter-patient heterogeneity during principal component analysis (e.g. using partial redundancy analysis) revealed consistent changes among cell types, with clear clustering of cell types and/or microenvironments based on metabolite profile (Fig. 2c). Analyses of single metabolites underscored these effects and revealed dramatic differences among cell types and microenvironments. Notably, the most extreme difference observed was for 1-methylnicotinamide (MNA), which was enriched in CD45-cells in general, and ∼10-100-fold in T cells when they infiltrated the tumor (Fig. 3a). This effect was most pronounced for CD4+ T cells; while MNA in CD8+ cells also appeared to be strongly affected by the environment, this was not significant as tumor CD8+ fractions were only evaluable for three of the six patients.

**Figure 2.**
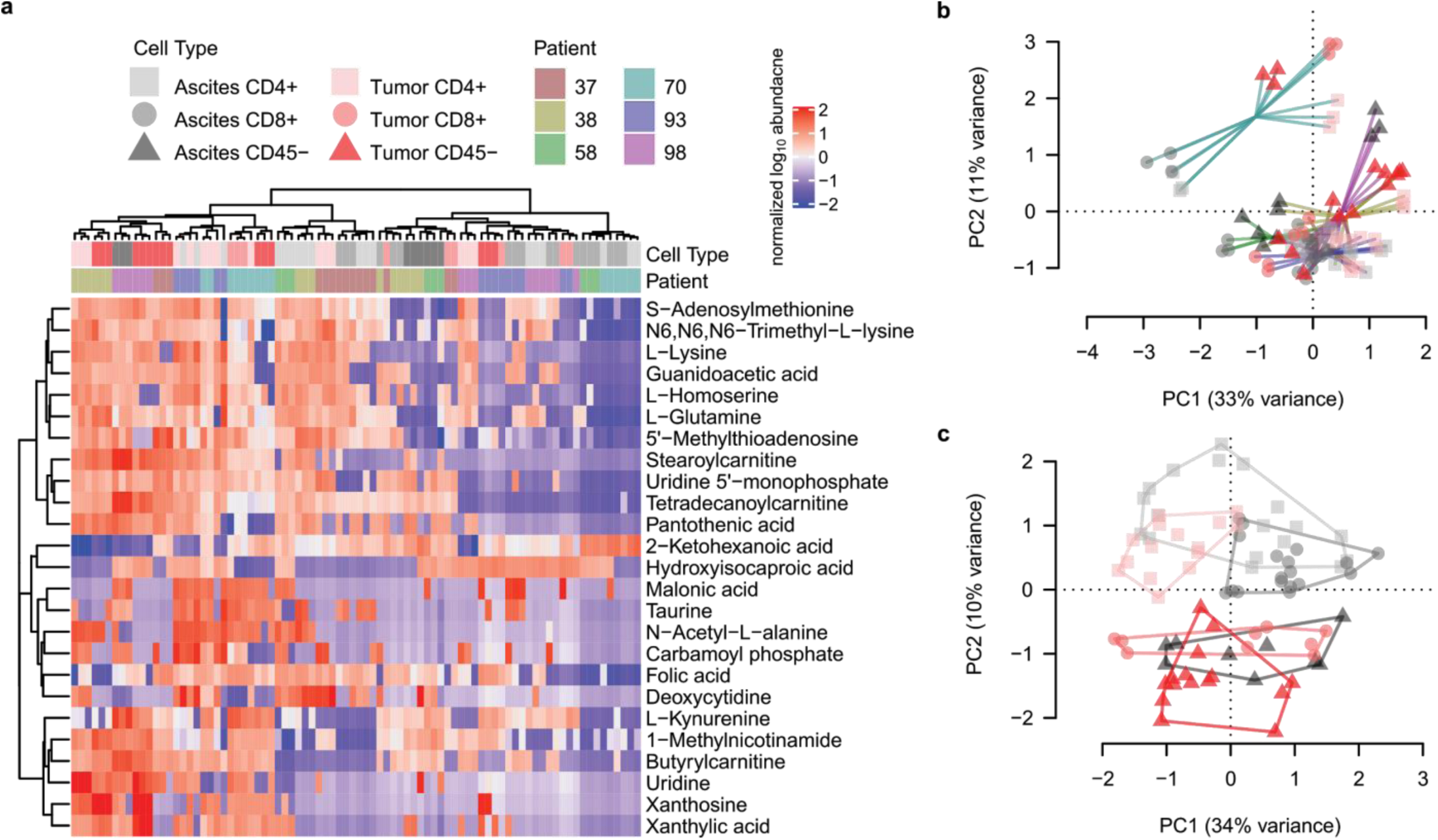
Metabolite profiling of matched ascites and tumor reveals key differences between tumor cells and T cells. **a**, Heatmap of normalized metabolite abundance, with dendrograms representing Ward’s clustering of Euclidean distances among samples. **b**, Principal components analysis (PCA) of sample metabolite profiles, showing triplicate replicates of each sample, with samples from the same patients joined by lines. **c**, PCA of sample metabolite profiles conditioned on patient (i.e. using partial redundancy); sample types are circumscribed by convex hulls.

**Figure 3.**
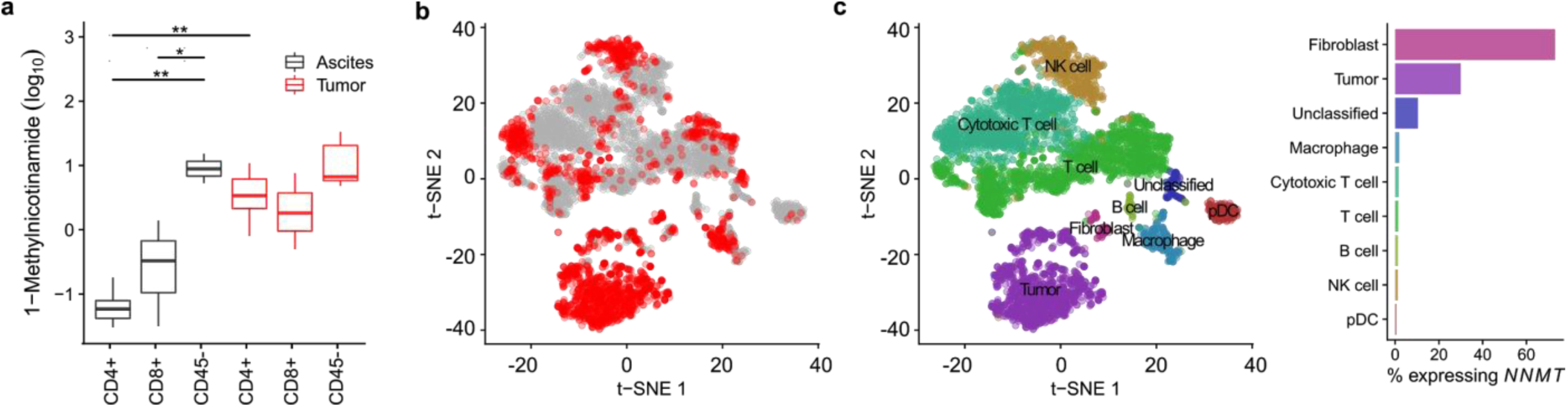
1-Methylnicotinamide (MNA) is more abundant in T cells from the tumor compared to ascites. **a**, Normalized abundance of MNA in CD4+, CD8+ and CD45-cells from ascites and tumor. Boxplots show medians (lines), interquartile range (box hinges) and range of data up to 1.5X interquartile range (box whiskers). P-values are determined using *limma* with patient as a random effect, as described in methods (*p<0.05, **p<0.01). **b**, t-SNE of scRNA-seq of ascites (grey) and tumor (red) (n=3 patients). **c**, *NNMT* expression in different cellular populations identified using scRNA-seq.

MNA is produced by the transfer of a methyl group from s-adenosyl-L-methionine (SAM) to nicotinamide (NA) by nicotinamide N-methyltransferase (NNMT). *NNMT* is over-expressed in multiple human cancers and has been linked to proliferation, invasion, and metastasis. To better understand the source of MNA in T cells in the TME, we used single cell RNA sequencing (scRNA-seq) to characterize *NNMT* expression across cell types in the ascites and tumor of three patients with HGSC (Extended Data Table 3). Profiling ∼6,500 cells revealed that *NNMT* expression was confined to presumptive fibroblast and tumor cell populations in both the ascites and tumor environments (Fig. 3b,c). Notably, there was no appreciable *NNMT* expression in any *PTPRC*-expressing (CD45+) populations (Fig. 3c), suggesting the MNA detected in metabolite profiling is imported into T cells. The expression of aldehyde oxidase 1 (*AOX1*), which converts MNA to 1-methyl-2-pyridone-5-carboxamide (2-PYR) or 1-methyl-4-pyridone-5-carboxamide (4-PYR), was likewise restricted to fibroblast populations (Extended Data Fig. 6), collectively suggesting that T cells lack the capacity for conventional MNA metabolism. This metabolite profile and scRNA-seq analysis also revealed similar, although less dramatic, patterns for both L-kynurenine and adenosine (Extended Data Fig. 7), two well-characterized immunosuppressive metabolites that were also elevated in T cells from the tumor, and/or in tumor cells. These trends, coupled with the striking enrichment of MNA in T cells within the tumor, raised the possibility that secretion of MNA into the TME may modulate the phenotypes of TIL to compromise antitumor immunity.

To determine the impact of MNA on T cells, healthy donor T cells were activated in the presence of MNA and assessed for proliferation and function. Addition of MNA did not lead to decreased proliferation or viability in either CD4+ or CD8+ T cells after 7 days (Fig. 4a), but rather increased the proportion of CD4+ and CD8+ T cells that expressed tumor necrosis factor alpha (TNFα) (Fig. 4b). While TNFα has been reported to have context-dependent pro- and anti-tumor effects, it has a well-described role in promoting ovarian cancer growth and metastasis^18–20^. Patients with ovarian cancer have been reported to have higher concentrations of TNFα within their ascites and tumor tissue than selected benign tissue^21–23^. Mechanistically, TNFα can modulate activation, function and proliferation of leukocytes, and change the phenotype of cancer cells^24,25^. Consistent with these findings, differential expression analysis of T cell populations between the ascites and tumor also revealed a significant up-regulation of *TNF* on T cells in the tumor relative to the ascites. Importantly, the increase in *TNF* was only apparent for T cell populations that did not exhibit a cytotoxic phenotype (Fig. 4c). Taken together, these data support the notion of a dual immune suppressive and tumor promoting role for MNA in HGSC.

**Figure 4.**
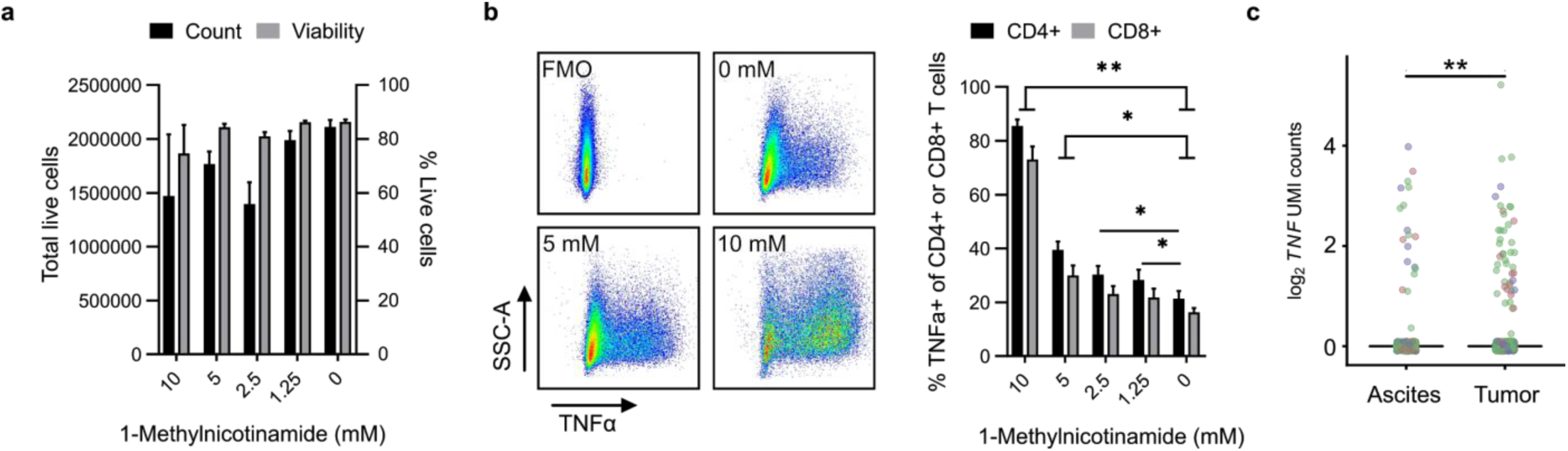
Exogenous MNA enhances TNFα expression in T cells. **a**, Total live cell count and viability directly from culture on day 7. Bar graphs represent mean with SEM of three healthy donors. **b**, TNFα expression in T cells treated with exogenous MNA. T cells were activated using CD3/CD28 with IL2 in respective concentrations of MNA for 7 days. Cells were stimulated with PMA/Ionomycin with GolgiStop™ for 4 hours prior to analysis. Example plot of live cells (left) and tabulated data (right). Bar graphs represent mean with SEM of 3 healthy donors. P-value determined using paired t-test (*p<0.05, **p<0.01). **c**, T cells (non-cytotoxic) show increased expression of *TNF* in the tumor relative to the ascites of HGSC. Colors represent different patients. Displayed cells have been randomly subsampled to 300 and jittered to limit overplotting (P_adj_ = 0.0076).

By applying a combined metabolomics approach, this study revealed pervasive immune metabolome differences between cells within the tumor and ascites of HGSC patients. This integrated analysis demonstrated differences in glucose uptake and mitochondrial activity between T cells and tumor cells in HGSC. However, these flow-based methods of assessing metabolism, while methodologically straightforward and providing single cell resolution, do not provide sufficient information regarding the cellular impacts of specific metabolites that function in cis or in trans within a given cell type. Importantly, our work uncovered a previously unrecognized metabolite MNA as differentially abundant between compartments and cell types. *In vitro*, MNA increased T cell-mediated secretion of the tumor promoting cytokine TNFα, providing insight for an alternative and non-cell autonomous role of MNA as an immune modulator in the ovarian TME.

## Methods

### Patient sample collection and processing

Patient specimens and clinical data were obtained through the BC Cancer Tumour Tissue Repository (TTR), certified by the Canadian Tissue Repository Network. All specimens and clinical data were obtained with either informed written consent or a formal waiver of consent under protocols approved by the Research Ethics Board of the BC Cancer Agency and the University of British Columbia (H07-00463). Samples are stored in a certified BioBank (BRC-00290). Detailed patient characteristics are shown in **Extended Data Table 1** and **Extended Data Table 3**.

For cryopreservation, patient tumor samples were mechanically disaggregated using a scalpel and pushed through a 100 μm filter to obtain a single cell suspension. Patient ascites was centrifuged at 1500 rpm for 10 minutes at 4 °C to pellet cells and remove supernatant. Cells obtained from tumor and ascites were cryopreserved in 50% heat inactivated human AB serum (Sigma), 40% RPMI-1640 (Fisher) and 10% DMSO.

### Cell culture reagents

Complete media consisted of a 0.22 μm filtered 50:50 supplemented RPMI1640:AimV. RPMI1640 + 2.05 mM L-Glutamine (Fisher) was supplemented with 10% heat inactivated human AB serum (Sigma), 12.5 mM HEPES (Fisher), 2 mM L-Glutamine (Fisher), 1x Penicillin Streptomycin solution (Fisher) and 50 μM B-mercaptoethanol. AimV (Invitrogen) was supplemented with 20 mM HEPES (Fisher) and 2 mM L-glutamine (Fisher). Flow cytometry staining buffer consisted of 0.22 μm filtered PBS (Invitrogen) supplemented with 3% heat inactivated AB human serum (Sigma). Cell enrichment buffer consisted of 0.22 μm filtered PBS supplemented with 0.5% heat inactivated human AB serum (Sigma).

### Flow cytometry for metabolic profiling

Cells were stained with 10 nM MitoTracker Deep Red (MT DR) and 100 μM 2-(N-(7-Nitrobenz-2-oxa-1,3-diazol-4-yl)Amino)-2-Deoxyglucose (2-NBDG) for 30 minutes in complete media at 37°C. Next, cells were stained with viability dye-eF506 for 15 minutes at 4 °C. Cells were resuspended in F_C_ block (eBioscience) and Brilliant Stain Buffer (BD Bioscience) diluted in flow cytometry staining buffer (according to manufacturer instructions) and incubated for 10 minutes at room temperature. Cells were stained with a panel of antibodies (**Extended Data Table 3**) in flow cytometry staining buffer for 20 minutes at 4 °C. Cells were resuspended in flow cytometry staining buffer prior to analysis (Cytek Aurora; 3L-16V-14B-8R configuration).

Cytometry data were analyzed using SpectroFlo and FlowJo V10, and figures were created using GraphPad Prism 8. Median Fluorescent Intensity of 2-NBDG and MT DR were log_10_ normalized prior to statistical analysis using paired t-test to account for matched patients. Any population with less than 40 events was removed from the analysis, an MFI value of 1 was inputted for any negative values prior to statistical analysis and data visualization.

### Unbiased discovery of cell populations in flow cytometry data

To supplement our manual gating strategy for the above flow panel, we used Full Annotation Using Shape-constrained Trees (FAUST)^14^ to automatically assign cells to populations, after dead cell exclusion in FlowJo. We manually curated outputs to merge populations that appeared to be mis-assigned (merged PD1+ with PD1-tumor cells), and retained populations comprising, on average, more than 2% of cells in each sample, for a total of 11 populations.

### Cell activation and enrichment for metabolite profiling optimization

Peripheral blood mononuclear cells (PBMCs) were isolated from a Leukopheresis Pack (Stemcell) using Ficoll gradient density centrifugation. CD8+ T cells were isolated from the PBMCs using CD8 MicroBeads (Miltenyi) and expanded using TransAct (Miltenyi) for 2 weeks in complete media according to manufacturer’s instructions. Cells were rested for 5 days in complete media with 10 ng/ml IL-7 (Peprotech) and then restimulated with TransAct. On day 7, cells were enriched using CD45 MicroBeads (Miltenyi) in three rounds of sequential enrichment according to the manufacturer’s instructions. Cells were aliquoted for analysis by flow cytometry (described above) and 1 million cells were aliquoted in triplicate for analysis by LC-MS/MS. Samples were processed for LC-MS/MS as described below. We imputed missing metabolite values with an ion count of 1000. Each sample was normalized by total ion count (TIC) and log_10_ transformed prior to statistical analysis.

### Cell-type enrichment of patient samples

Patient cells were filtered through a 40 μm filter. Samples were enriched for CD8+, CD4+ and CD45-cells (on ice) using three sequential rounds of positive selection by magnetic bead separation (Miltenyi MACS MicroBeads). CD8-fraction was used for CD4 enrichment, and the CD4-fraction was used for CD45-enrichment to maximize cell recovery.

### LC-MS/MS metabolite profiling

To prepare samples (in triplicate) for metabolite profiling, cells were washed once with ice-cold saline solution and 1 mL of 80% methanol added to each sample before vortexing and snap freezing in liquid nitrogen. Samples were subjected to 3 freeze-thaw cycles, and centrifuged at 14,000 rpm for 15 minutes at 4 °C. The metabolite-containing supernatant was evaporated until dry. Metabolites were reconstituted in 50 μl of 0.03% formic acid, vortex-mixed, and centrifuged to remove debris. The supernatant was transferred to a high-performance liquid chromatography (HPLC) vial for the metabolomics study. Each sample was processed with similar numbers of cells using a randomized processing scheme to prevent batch effects. We performed qualitative assessment of global metabolites as previously published on the AB SCIEX QTRAP 5500 triple-quadrupole mass spectrometer^**26**^. Chromatogram review and peak area integration were performed using MultiQuant software version 2.1 (Applied Biosystems SCIEX).

### Characterizing metabolic differences across cell types and microenvironments

Missing metabolite values were imputed with an ion count of 1000 and normalized peak area calculated for each detected metabolite using the total ion count from each sample to correct variations introduced from sample handling through instrument analysis. TIC-normalization was followed by log_10_ transformation and autoNorm row-scaling using *MetaboAnalystR*^**27**^ (default parameters). We conducted exploratory analysis of metabolome differences across sample types using PCA with the *vegan* R package, and conditioned the analysis on patient using partial redundancy analysis. Heatmap dendrograms were constructed using Ward’s method to cluster Euclidean distances among samples. We identified differentially-abundant metabolites across cell types and microenvironments using *limma*^**28**^ on the log_10_-transformed row-normalized metabolite abundances. To simplify interpretation, we specified the model using the group means parameterization, treating cell types within microenvironments as each group (n=6 groups); for significance testing we took the average of triplicate measurements for each metabolite to avoid pseudoreplication, and included patient as a block in the *limma* design. To examine metabolites that differed across patients we re-fit models in *limma* including patient as a fixed effect. We reported significance at P_adj_ < 0.05 (Benjamini-Hochberg correction) for pre-specified contrasts among cell types and microenvironments.

### scRNA-seq

Single cell transcriptome sequencing was performed on total viably-frozen ascites and tumor samples using the 10X 5’ Gene Expression protocol, following viability enrichment with the Miltenyi Dead Cell Removal Kit (>80% viability). 5 cases with matched tumor and ascites available were profiled, although low viability from 1 tumor sample prevented its inclusion. To enable multiplexing of patients, we combined samples from each patient in lanes of the 10X Chromium controller, with separate runs for ascites and tumor fractions. Following sequencing (Illumina HiSeq 4000 26×98bp PE, Genome Quebec; mean of 73,488 and 41,378 reads per cell for tumor and ascites, respectively), we assigned donor identities using *CellSNP* and *Vireo*^**29**^ (based on the common human SNP VCF provided by *CellSNP* for GRCh38). We excluded unassigned cells and those identified as doublets, and matched donors between ascites and tumor samples based on the nearest identity-by-state (IBS) of inferred patient genotypes using *SNPRelate*^**30**^. Based on this assignment, we retained 3 cases with abundant cellular representation in both tumor and ascites fractions for downstream analysis. Following quality filtering steps in the *scater*^**31**^ and *scran*^**32**^ BioConductor packages, this yielded 6,975 cells (2,792 and 4,183 from tumor and ascites, respectively) for analysis. We clustered cells by expression using *igraph*’s^**33**^ Louvain clustering implementation of the shared nearest neighbour network (SNN) based on Jaccard distance. Clusters were manually annotated into presumptive cell types based on marker gene expression and visualized with t-SNE.

### T cell functional assay

PBMCs were isolated from a leukapheresis product (Stemcell) by Ficoll gradient density centrifugation. CD3+ cells were isolated from the PBMCs using CD3 Microbeads (Miltenyi). The CD3+ cells were activated with plate bound CD3 (5 μg/ml), soluble CD28 (3 μg/ml), and IL-2 (300 U/ml, Proleukin) in the presence or absence of MNA. On the final day of expansion viability (Fixable Viability Dye eFluor450, eBioscience) and proliferation (123count eBeads, Thermo) was assessed by flow cytometry. Effector function was assessed by stimulating cells for 4 hours with PMA (20 ng/ml) and Ionomycin (1 μg/ml) with GolgiStopTM and monitored for CD8-PerCP (RPA-T8, Biolegend), CD4-AF700 (RPA-T4, Biolegend), and TNFa-FITC (MAb11, BD).

### Statistical analysis

Statistical analysis was carried out as described in the text or methods using GraphPad Prism 8, Microsoft Excel or R v3.6.0. Where multiple samples were taken from the same patient (e.g. ascites and tumor), we used paired t-tests, or included patient as a random effect in linear or generalized models, as appropriate. For metabolomic analysis, significance testing was done on means of triplicate measurements.

## Data Availability

Raw sequencing data will be deposited at NCBI dbGAP (Accession pending). Processed data files and scripts to reproduce metabolomics and scRNA-seq analyses are available at github.com/vicDRC/onecarbon. Flow cytometry data will be deposited at flowrepository.

## Code Availability

R scripts to reproduce metabolomic and single cell RNA-seq analyses are available at github.com/vicDRC/onecarbon.

## Acknowledgements

Thank you to the patients who donated specimens to make this study possible. We thank Mary Lesperance and Farouk Nathoo for advice on statistical analysis. This work was supported by the research grants to J.J.L from the US Department of Defence Ovarian Cancer Research Program Pilot Award (W81XWH-18-1-0264), Canadian Institutes for Health Research (MOP-142351, PJT-162279), to P.T.H from the Carraresi Family Foundation Award, to B.H.N from the BC Cancer Foundation, and the Canadian Foundation for Innovation. M.K.K is supported by a University of Victoria Graduate Award. P.T.H. is supported by a Canadian Institutes for Health Research Post-Doctoral Fellowship. Biospecimens were obtained with assistance from the IROC-TTR biobank. Single cell library preparations and RNA sequencing were provided by Genome Quebec.

## Author Contributions

M.K.K. and P.T.H. designed and performed experiments, analyzed the data, and wrote the manuscript. S.M., S.K., and B.P. helped perform the metabolite profiling experiments. P.T.H., J.S, B.A., J.S, and B.H.N. generated the scRNA-seq data. L.Z. and R.J.D. generated and analyzed the LC-MS/MS data. P.H.W. oversaw biospecimen collection through the IROC-TTR. J.J.L. conceived the project and wrote the manuscript.

## Competing Interests Statement

J.S. is a permanent member and owns stocks of Surface Oncology. R.J.D is a member of the Scientific Advisory Board at Agios Pharmaceuticals.

## Corresponding Author

Correspondence to Julian J. Lum.

## Extended data

**Extended Data Figure 1.**
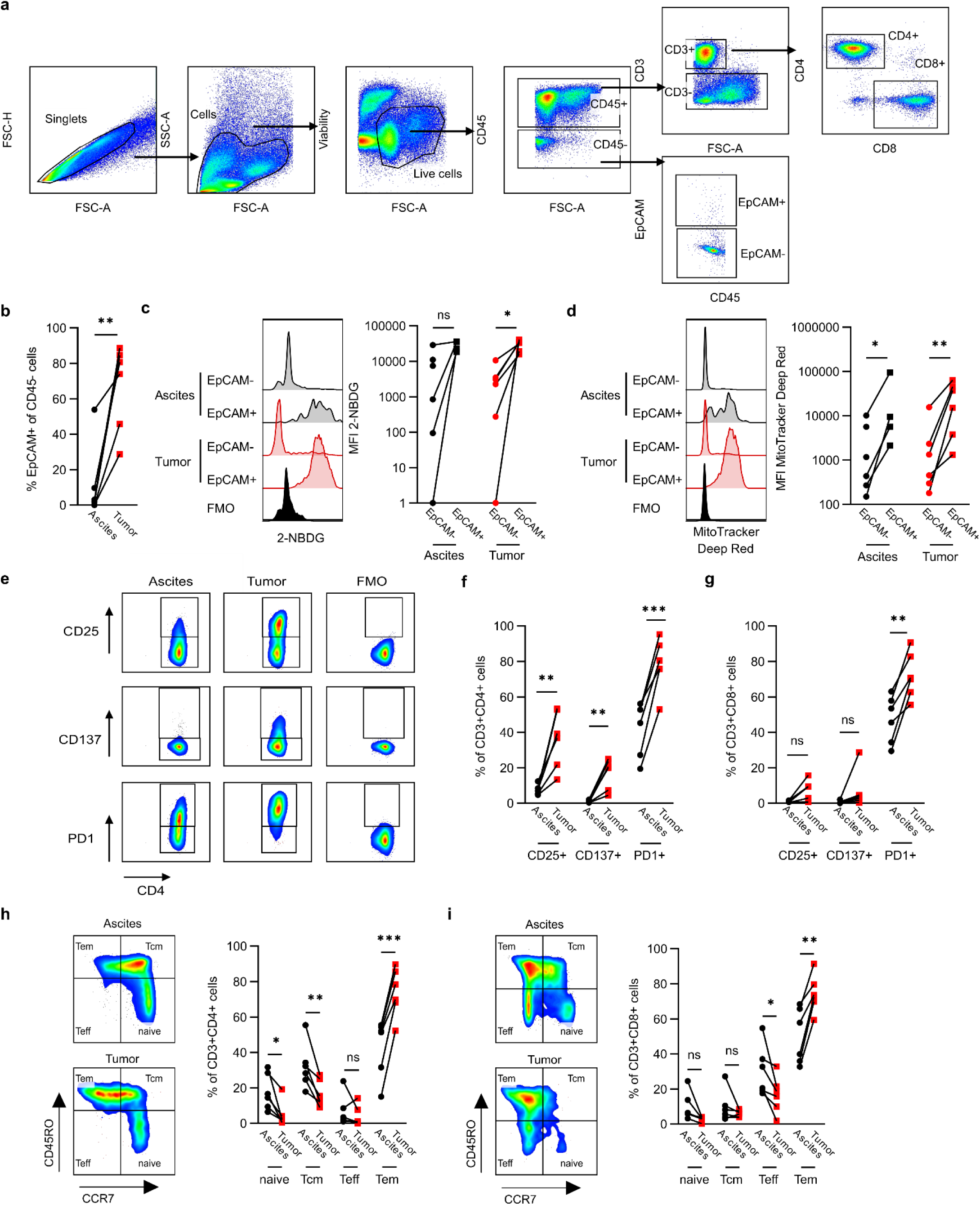
Phenotypic characterization of ascites and tumor by flow cytometry. **a**, Representative gating strategy for analysis by flow cytometry. **b**, Proportion of EpCAM+ (of CD45-) tumor cells within ascites and tumor. **c, d**, Representative plot (left) and tabulated data (right) for glucose uptake (2-NBDG) (**c**) and mitochondrial activity (MitoTracker Deep Red) (**d**) of EpCAM+CD45-tumor and EpCAM-CD45-stromal cells from ascites and tumor. **e**, Representative gating strategy for CD25, CD137 and PD1 expression by flow cytometry. **f, g**, CD25, CD137 and PD1 expression on CD4+ T cells (**f**) and CD8+ T cells (**g**). **h, i**, Naive, central memory (Tcm), effector (Teff), and effector memory (Tem) phenotype based on CCR7 and CD45RO expression. Representative plot (left) and tabulated data (right) for CD4+ T cells (**h**) and CD8+ T cells (**i**) from ascites and tumor. P-values (*p<0.05, **p<0.01, ***p<0.001) determined by paired t-test (**b-d, f-i**). Median Fluorescence Intensity (MFI).

**Extended Data Figure 2.**
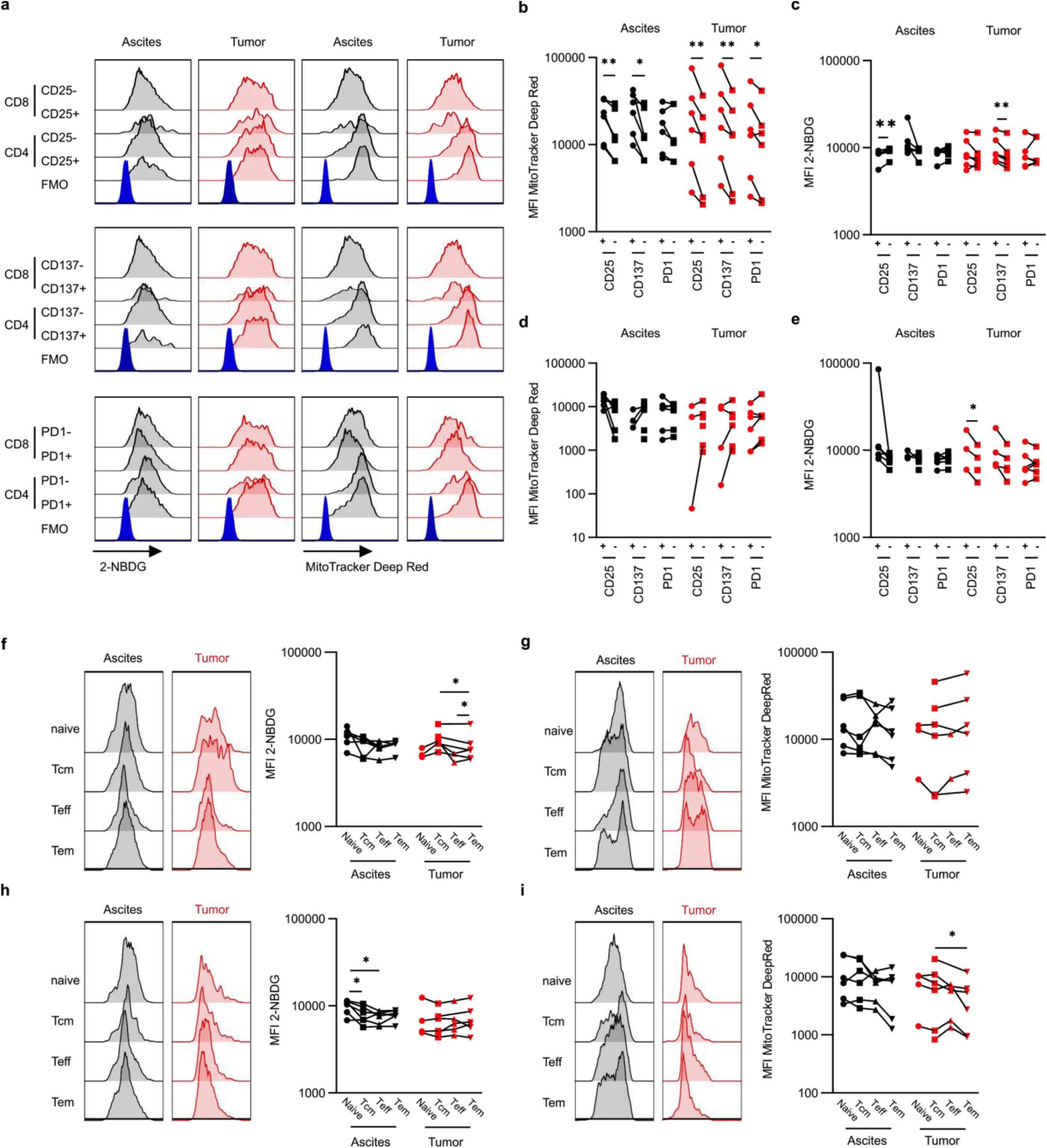
T cell metabolism is impacted by expression of activation markers. **a**, Representative plots of glucose uptake (2-NBDG) and mitochondrial activity (MitoTracker Deep Red) for CD25, CD137 and PD1 positive and negative CD4 and CD8 T cells. **b, c**, Mitochondrial activity (**b**) and glucose uptake (**c**) of CD25, CD137 and PD1 CD4+ T cells. **d, e**, Mitochondrial activity (**d**) and glucose uptake (**e**) of CD25, CD137 and PD1 CD8+ T cells. **f, g**, Representative plot (left) and tabulated data (right) for glucose uptake (**f**) and mitochondrial activity (**g**) of naive, Tcm, Teff and Tem CD4+ T cells. **h, i**, Representative plot (left) and tabulated data (right) for glucose uptake (**h**) and mitochondrial activity (**i**) of naive, Tcm, Teff and Tem CD8+ T cells. P-values (*p<0.05, **p<0.01, ***p<0.001) determined by paired t-test **b-i**). Lines indicate matched patients (**b-i**). Fluorescence Minus One (FMO); Median Fluorescence Intensity (MFI); Central memory T cells (Tcm); Effector T cells (Teff); Effector memory T cells (Tem).

**Extended Data Figure 3.**
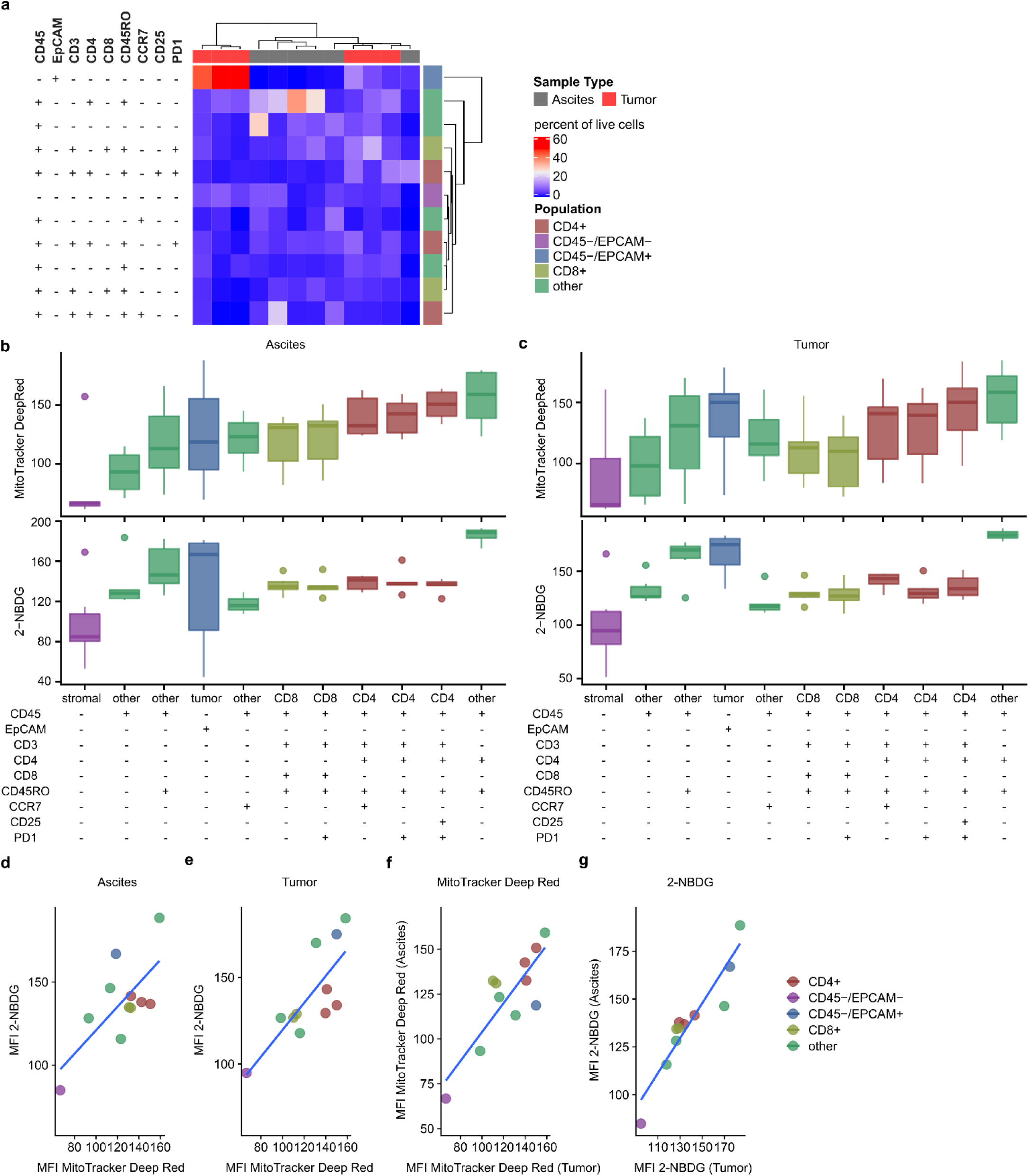
Automated analysis of metabolism and cell type by flow cytometry. **a**, Heat map of cell type abundance. **b, c**, Glucose uptake and mitochondrial activity of cell fractions within ascites (**b**) and tumor (**c**). Boxplots show medians (lines), interquartile range (box hinges) and range of data (box whiskers; excepting outliers, shown as points) **d, e**, Correlation between glucose uptake and mitochondrial activity of cell fractions within the ascites (**d**) and tumor (**e**). **f, g**, Correlation of glucose uptake (**f**) and mitochondrial activity (**g**) of cell fractions between ascites and tumor. Median (per phenotype) of median (per sample) biexponential-transformed Median Fluorescence Intensity (MFI) values shown.

**Extended Data Figure 4.**
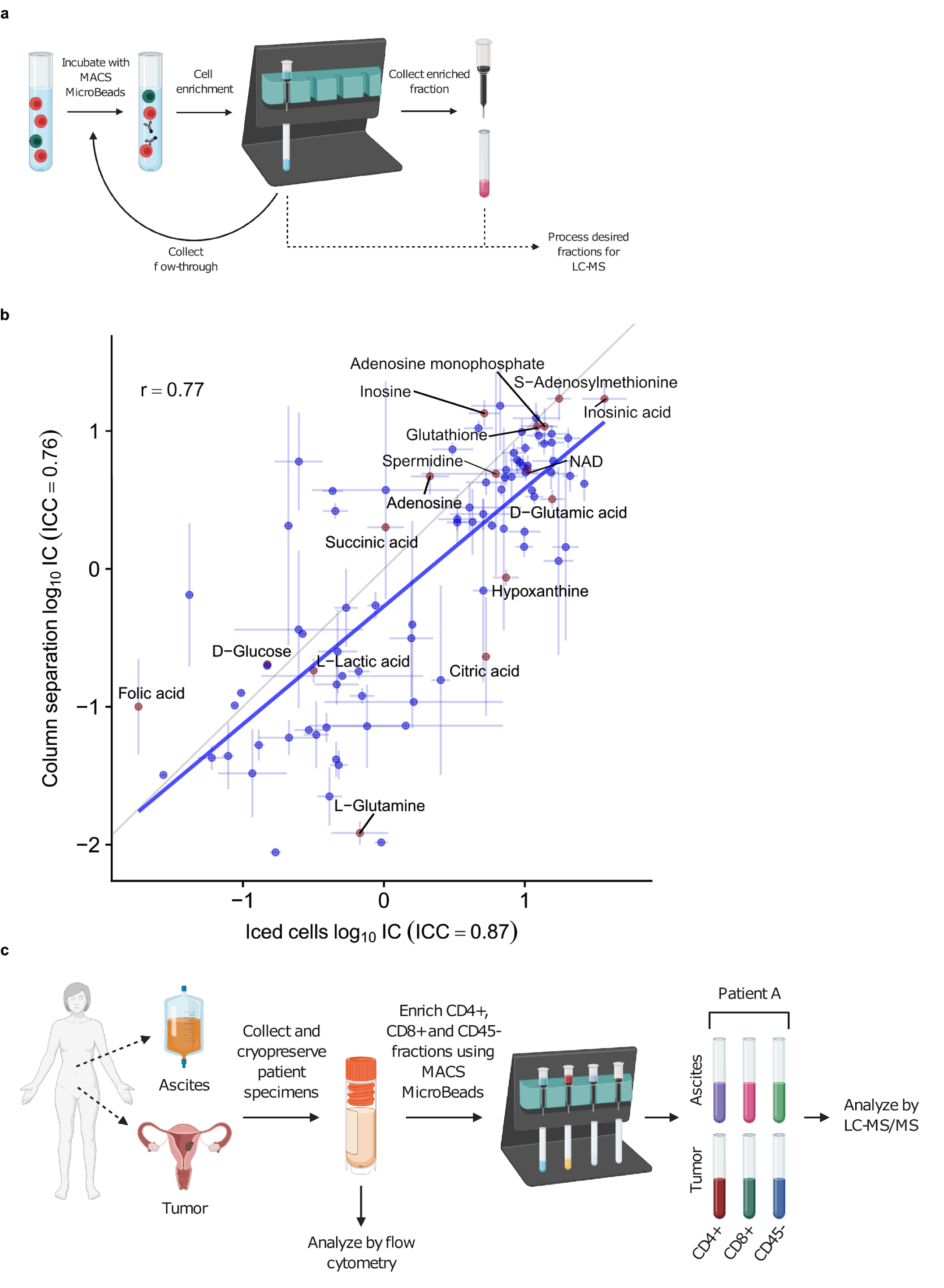
Schematic workflow and impact of enrichment on metabolite profiling. **a**, Schematic of magnetic bead enrichment. Cells underwent three consecutive rounds of magnetic bead enrichment or remained on ice. **b**, Impact of enrichment type on metabolite abundance. Means of triplicate measurements for each enrichment type +/- SE shown. Gray line represents 1:1 relationship. Intraclass correlation for replicate measurements (ICC) shown in axis labels. **c**, Schematic of patient metabolite profiling workflow. Ascites or tumor was collected from patients and cryopreserved. A fraction of each sample was analyzed by flow cytometry, while the remaining sample underwent three rounds of enrichment for CD4+, CD8+ and CD45-cells. These cell fractions were analyzed using LC-MS/MS.

**Extended Data Figure 5.**
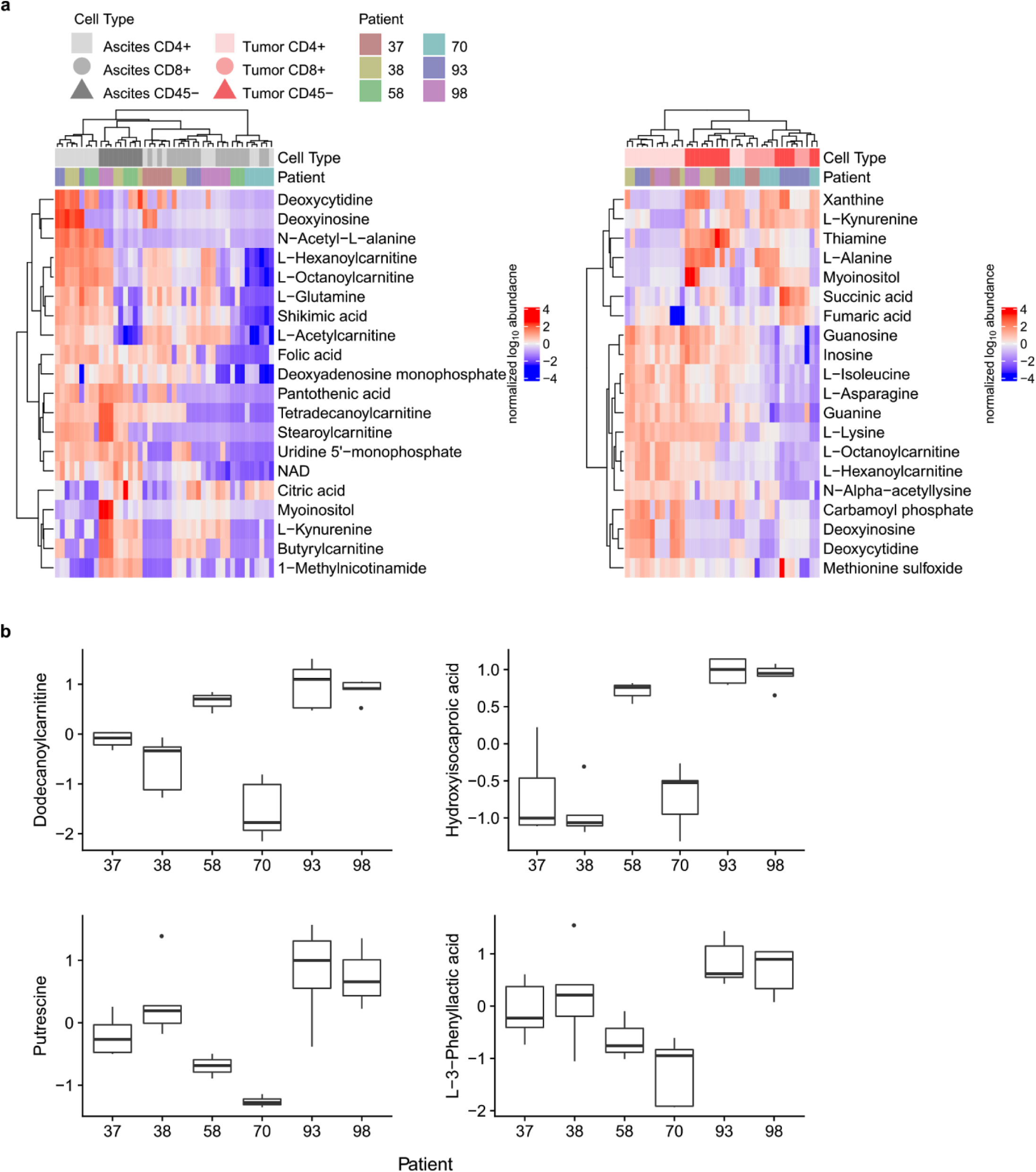
Changes in relative metabolite abundance across cell types within ascites and tumor. **a**, Heatmap of normalized metabolite abundance, with dendrograms representing Ward’s clustering of Euclidean distances among samples. Relative abundance of metabolites in the ascites (left) and tumor (right). **b**, Top four significantly differing metabolites across patients (all P_adj_ for F-Test of patient effect in *limma* < 0.05). Boxplots show medians (lines), interquartile range (box hinges) range of data up to 1.5X interquartile range (box whiskers; excepting outliers, shown as points).

**Extended Data Figure 6.**
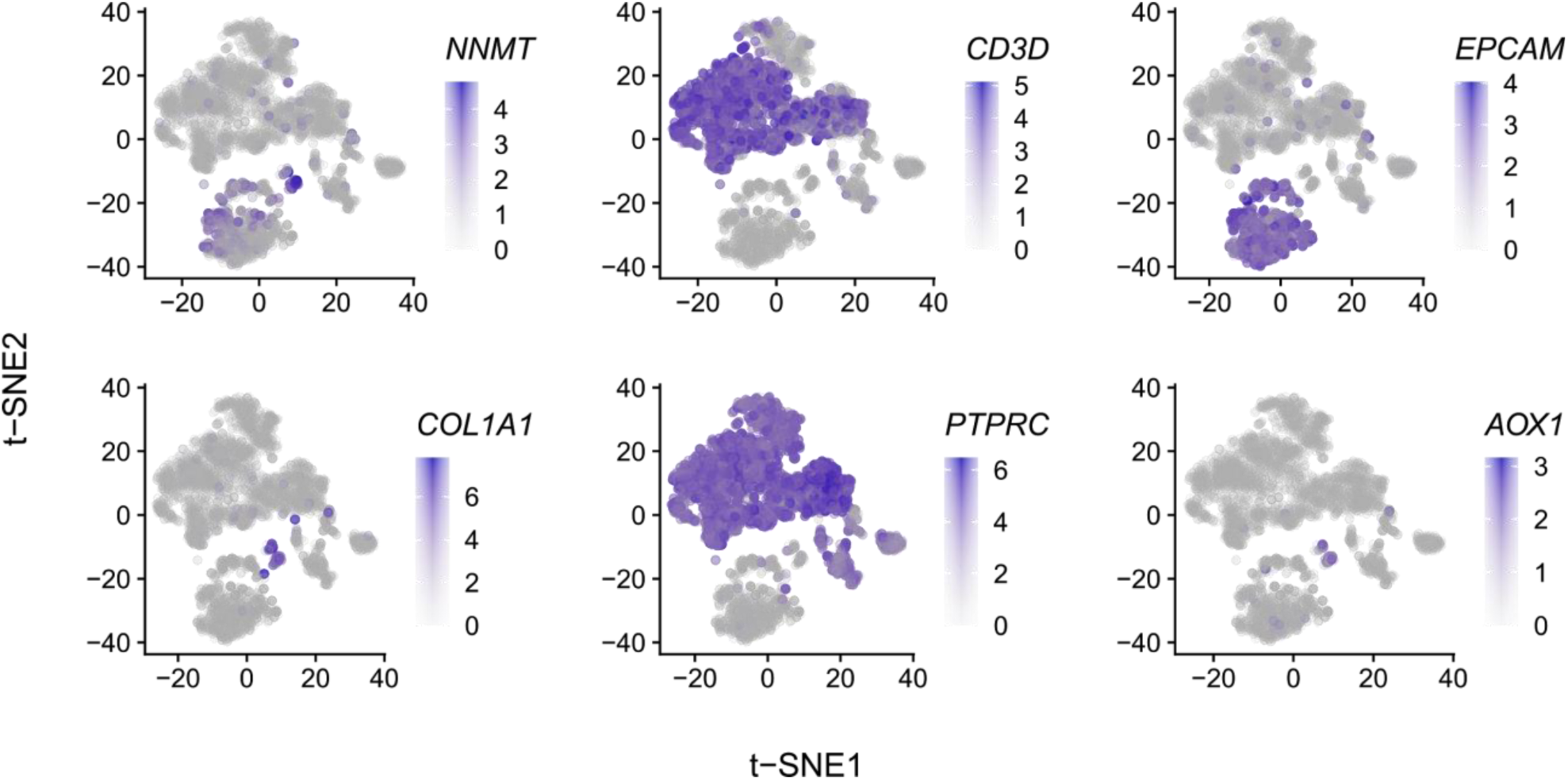
Expression of population defining makers and metabolic genes within the scRNA-seq data. Expression of *NNMT, CD3D, EPCAM, COL1A1, PTPRC* and *AOX1* within ascites and tumor, shown as log_2_ normalized unique molecular identifier (UMI) counts.

**Extended Data Figure 7.**
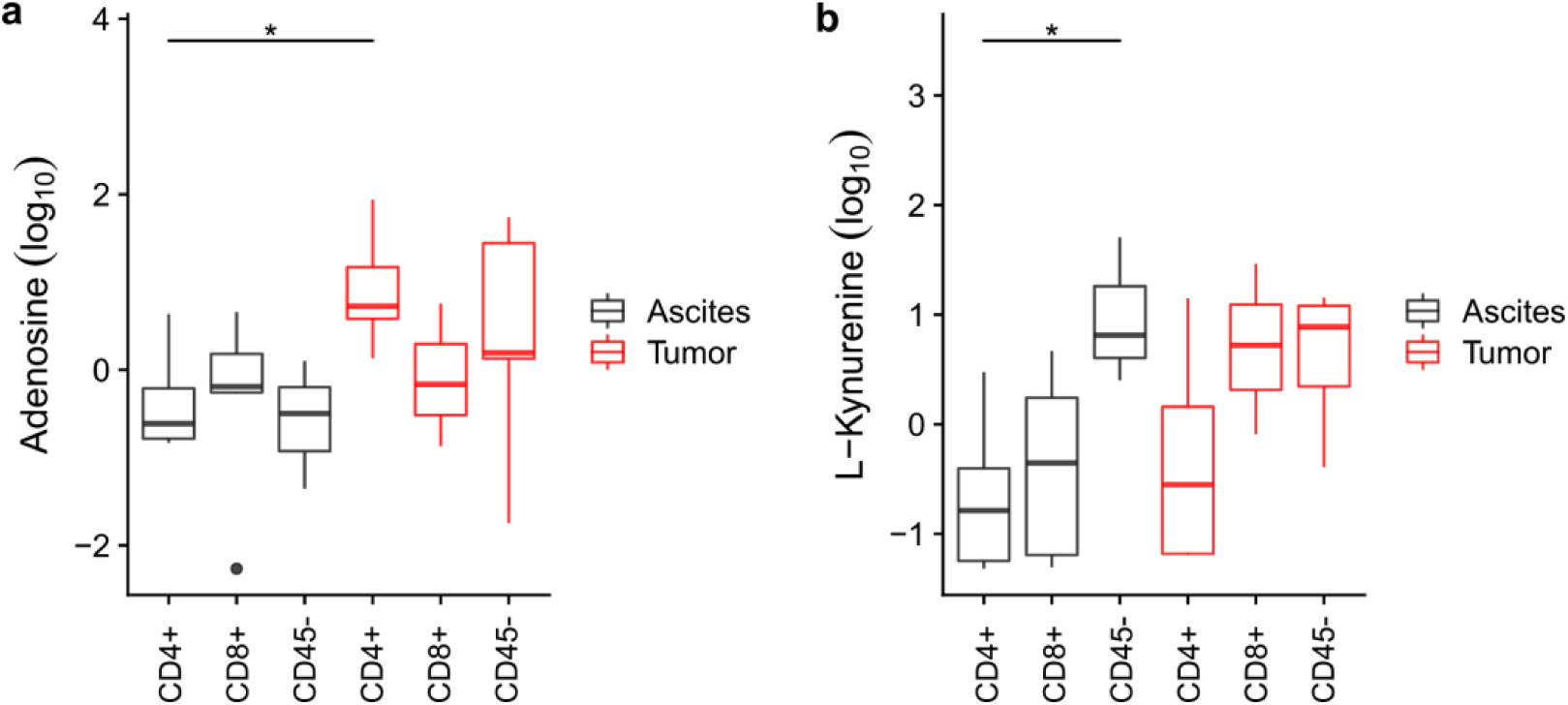
Relative abundance of adenosine (a) and L-kynurenine (b) measured by LC-MS/MS. P-values determined as described in methods (*p<0.05). Boxplots show medians (lines), interquartile range (box hinges) and range of data up to 1.5X interquartile range (box whiskers; outliers shown as points). P values determined using *limma* as per Figure 3.

**Extended Data Table 1.** HGSC patient characteristics for metabolic profiling by flow cytometry and LC-MS/MS.

| Patient | Tumor location | Age at surgery | Survival (months)* | Tumor stage<sup>†</sup> | Tumor grade<sup>†</sup> | Ascites volume (mL) |
| --- | --- | --- | --- | --- | --- | --- |
| 37 | omentum | 70 | 12 | 3C | 3 | 1800 |
| 38 | ovary | 40 | 16 | T3b | 3 | 1200 |
| 70 | omentum | 50 | 23 | 3C | 3 | 80 |
| 58 | omentum | 78 | 18 | 4 | 3 | 2000 |
| 93 | fallopian tube | 53 | 82 | 3B | 2 | 1600 |
| 98 | omentum | 39 | 13 | 3C | 3 | 2000 |

\* Survival calculated from the date of diagnosis to the date of death
<sup>†</sup> Tumor stage and grade were determined by a pathologist at the time of surgery

**Extended Data Table 2.** Flow cytometry metabolic profiling panel.

| Fluorochrome | Marker | Expression | Clone | Company | Catalogue number |
| --- | --- | --- | --- | --- | --- |
| --- | 2-NBDG | Glucose uptake | --- | Thermo | N13195 |
| PE | CD326 (EpCAM) | Epithelial cells | 1B7 | Thermo | 12-9326-42 |
| PerCP | CD8 | Effector T cells | RPA-T8 | Biolegend | 301030 |
| PerCP-eFluor710 | CD25 | Activation/Tregs | 4E3 | Thermo | 46-0257-41 |
| PE-Cy7 | CD45RO | Phenotype | UCHL1 | Thermo | 25-0457-42 |
| --- | MitoTracker Deep Red | Mitochondrial activity | --- | Thermo | M22426 |
| AF700 | CD4 | Helper T cells | RPA-T4 | Biolegend | 300526 |
| APC/Fire750 | CCR7 | Phenotype | G043H7 | Biolegend | 353246 |
| eFlour506 | Viability | Live/dead cells | --- | Thermo | 65-0866-14 |
| PO | CD45 | Leukocytes | HI30 | Thermo | MHCD4530 |
| BV605 | CD137 | Activation | 4B4-1 | Biolegend | 309822 |
| BV650 | CD279 (PD1) | Activation/Exhaustion | EH12.2H7 | Biolegend | 329950 |
| BV750 | CD3 | T cells | SK7 | Biolegend | 344845 |

**Extended Data Table 3.** HGSC patient characteristics for single cell RNA sequencing.

| Patient | Tumor location | Age at surgery | Survival (months)* | Tumor stage<sup>†</sup> | Tumor grade<sup>†</sup> |
| --- | --- | --- | --- | --- | --- |
| 46 | ovary | 54 | 24 | 3C | 3 |
| 58 | omentum | 78 | 18 | 4 | 3 |
| 59 | omentum | 68 | 26 | 3C | 3 |
| 69 | omentum | 59 | 17 | 3C | 3 |
| 109 | ovary | 77 | 62 | 3C | 3 |

\* Survival calculated from the date of diagnosis to the date of death
<sup>†</sup> Tumor stage and grade were determined by a pathologist at the time of surgery

